# Harnessing the Power of Sex Differences: What a Difference Ten Years Did Not Make

**DOI:** 10.1101/2021.06.30.450396

**Authors:** Rebecca K. Rechlin, Tallinn F.L. Splinter, Travis E. Hodges, Arianne Y. Albert, Liisa A.M. Galea

**Affiliations:** Department of Psychology, University of British Columbia, Vancouver, BC, Canada; Department of Biology, University of British Columbia, Vancouver, BC, Canada; Women’s Health Research Institute of British Columbia, University of British Columbia, Vancouver, BC, Canada; Djavad Mowafaghian Centre for Brain Health, University of British Columbia, Vancouver, BC, Canada; Women’s Health Research Cluster, University of British Columbia, Vancouver, BC, Canada

## Abstract

Sex differences exist in many neurological and psychiatric diseases. Mandates have been initiated across funding agencies for research to include males and females. What has been lacking in the literature is a detailed assessment of how sex is incorporated into the design (e.g. balanced design) and into the analyses (e.g. covariate). We surveyed papers in 2009 and 2019 across six journals in Neuroscience and Psychiatry. There was a 30% increase in the percentage of papers that included both sexes to 68% in 2019. Despite this increase, in 2019 only 19% of studies used an optimal design for discovery of possible sex differences and only 5% analyzed sex as a discovery variable. Here we show that little progress has been made in harnessing the power that sex differences can afford in research for discovery and therapeutic potential for neurological and psychiatric disease to improve the health of men, women and gender diverse individuals.

**Highlights:**

68% of Neuroscience and Psychiatry papers reported the use of both sexes in 2019
Only 19% of studies in 2019 used sex consistently throughout the study analyses
Of the studies that used males and females, 59% did not include sex in the analyses
Only 5% of studies in 2019 used sex as a discovery variable in their analyses
Male only papers were 8.4 times more prevalent than female-only papers

## Introduction

The consideration of sex in published reports is essential to our understanding of disease and the biological mechanisms that contribute to the etiology, manifestation and treatment of disease^1^. The study of sex differences is critical to our understanding of precision medicine in finding effective treatments for disease. Sex differences exist in the prevalence and manifestation of a number of neurological and psychiatric diseases^2,3^. Females are more likely to be diagnosed with multiple sclerosis, major depressive disorder, and have a greater lifetime risk of Alzheimer’s Disease compared to males, whereas males are more likely to be diagnosed with autism spectrum disorder, attention and hyperactivity disorder, and Parkinson’s Disease^1–4^. Even in diseases that do not show strong sex differences in prevalence, age of disease onset or manifestation can be different between the sexes^5,6^. Perhaps more concerning, there are notable differences in time to diagnosis^7^, disease progression^2,4^, vaccine response^8^ and treatment efficacy/drug response^9^. Harnessing the knowledge that males and females can differ on several disease-related outcomes will be fruitful in not only understanding disease but also in determining whether sex-specific risk factors for disease may warrant further attention. For example, the manifestation of cardiovascular disease can be different between the sexes, prompting calls for changes to the diagnostic guidelines for cardiovascular disease based on sex^10^. To make headway for precision medicine and most effective treatment and diagnoses, sex must be taken into consideration in the design and analyses of data.

Many health disparities in treatment and diagnosis have been attributed to the lack of research in females and inclusion of women in clinical trials^11,12^. To increase the enrolment of women in clinical research, the United States Congress passed The Revitalization Act of 1993. This Act stated that women and minorities must be included as subjects in clinical trials funded by the National Institutes of Health (NIH). However, implementation of the requirement of women and minorities has not translated into analysis by sex or race/ethnicity^13^. The importance of sex consideration in research led the NIH to further mandate the inclusion of women and minorities in clinical research in 2001, and finally the addition of sex as a biological variable (SABV) in biomedical research in 2016^14^. However, this mandate, much like the one for clinical trials in 1993, did not include specifications as to the analysis of the data by sex^15^ nor did it specify sample size requirements^16^. Other countries have notable differences in their recommendations, timeline and mandates. The Canadian Institutes of Health Research (CIHR) implemented Sex and Gender-Based Analysis (SGBA) in 2010 as a mandatory component and in 2019 into the scoring of grant. Horizon Europe (European Commission) has been working on policy changes since 2002 requiring the integration of sex and gender in research where relevant^17^ and in 2020 Horizon Europe has indicated the need for inclusive intersectionality analyses of gender and sex in 2020 (Supplement Figure 1). Although prescriptive guidelines from funding agencies are lacking there are a number of reviews with suggestions on the appropriate incorporation of SABV and SGBA in the literature^18–20^. Despite the mandates and recommendations there have been implementation issues of the mandate as reviewers and authors of papers may be applying SABV and SGBA inconsistently perhaps given the lack of official guidelines^21,22^.

The biomedical and clinical research community is beginning to make corrections for a long-standing bias of using males predominately in research. With the publication by Beery and Zucker^23^ on the lack of sex inclusion in the literature in publications from 2009 it became clear that, although there was considerable variation by research field, the majority of studies were not using both sexes^23^. Studies in human populations were more likely to use both males and females across the ten disciplines examined compared to studies using animals^23^. A ten-year follow up was done demonstrating a 29% increase from 2009 in the inclusion of both sexes in research to 49% of articles in 2019, with Neuroscience having one of the largest increases in sex inclusion^24^. Even though a greater proportion of studies are including both sexes, there are issues in how these sexes have been included, as approximately one third of sex-inclusive studies did not specify the sample size^24^ and the large majority of studies that used males and females failed to analyze the data by sex in 2009^23^. Furthermore, there was an 8% decrease in the papers that used sex in their analyses^24^ over the years as only one discipline (Pharmacology) improved in analyses of sex across the ten years. Furthermore, sex bias favouring males is still prevalent in neuroscience research^24,25^. In fact a study from Will and colleagues^25^ indicated that the use of solely males in studies increased from 2010 to 2014, whereas the number of female studies remained at a constant low value (5% in Neuroscience). Thus, across the 10 years, studies indicate that although the sex omission rate is decreasing across disciplines, the use of sex in the analyses and the large differential in single-sex studies favoring males have not appreciably changed^24^.

What has been lacking in the literature is a detailed assessment of how sex is reported in papers (whether the study design is balanced, sex used consistently throughout the studies within the papers) and how males and females are included in any analyses. Often in clinical studies, sex is used as a covariate which controls for sex by removing the linear variation due to sex from the analysis and does not inform on the effect of sex. Therefore, in the present study, we examined not only whether a statistical analysis was done in the studies but what type of analysis was done to determine whether sex was controlled for, via a covariate analyses, or explicitly examined as a discovery variable. We were also interested in how many papers used an experimental design that was optimal for discovery of potential sex differences (reporting sample size, relatively balanced design, sex used through throughout the experiments). We examined experimental design as an indication if the papers were addressing the possibility of noting sex differences in their data, with the understanding that not all papers would be designed to address sex differences.

Given the prominent sex differences in neurological and psychiatry disorders, we chose to do a detailed examination of journals that targeted Neuroscience and Psychiatry. As the mandates for inclusion of males and females in biomedical research were in place in 2016, we examined two years over the ten-year period of 2009 to 2019 as was done by Woitowich and colleagues^24^ and as these were dates before and after the recommendations from Horizon Europe, CIHR and NIH. We hypothesized that there would be an increase in the number of papers that included both sexes from 2009 to 2019 in Neuroscience and Psychiatry papers, but also that there would be an increase in experimental design that was not optimized to examine sex as a biological variable. We also expected that most studies that analyzed sex as a factor would do so without using sex as a primary discovery variable across both disciplines, irrespective of year. Here we show that although the vast majority of papers include both sexes, only 19% include an optimal design for the discovery of possible sexes and only 5% included sex as a discovery variable in 2019.

## Methods

We exhaustively examined research papers within three journals in Neuroscience and Psychiatry across two years. We chose journals based on the high ISI Clarivate rankings that published primary research papers. Three Neuroscience journals (*Nature Neuroscience*, *Neuron*, *Journal of Neuroscience*) and three Psychiatry journals (*Molecular Psychiatry*, *Biological Psychiatry*, *Neuropsychopharmacology*) were chosen. We assessed papers published in the year 2009 and in 2019 to assess whether there has been an increase in the inclusion of sexes, improvements to experimental design and analyses to examine potential differences between the sexes.

### Studies included

All primary research articles from 2009 and 2019 were analyzed if the papers used rats, mice, human subjects, or if fetal cells/cell lines were included. Cell lines included immortalised cell lines, primary cell culture, and stem cell derivatives. As sex of cells matters in a variety of outcomes^26^ these studies were included. Brief communications, reviews, viewpoints etc. were excluded. This resulted in a review of a total of 3191 publications (Figure 1). Assessments were done by two trained curators who had >99% interrater reliability (RKR, TFLS). When the categorization of analyses within the paper (see below) was questioned, these were confirmed by AYA, a biostatistician - who was consulted on 0.5% of the papers reviewed or 16 times in total.

**Figure 1.**
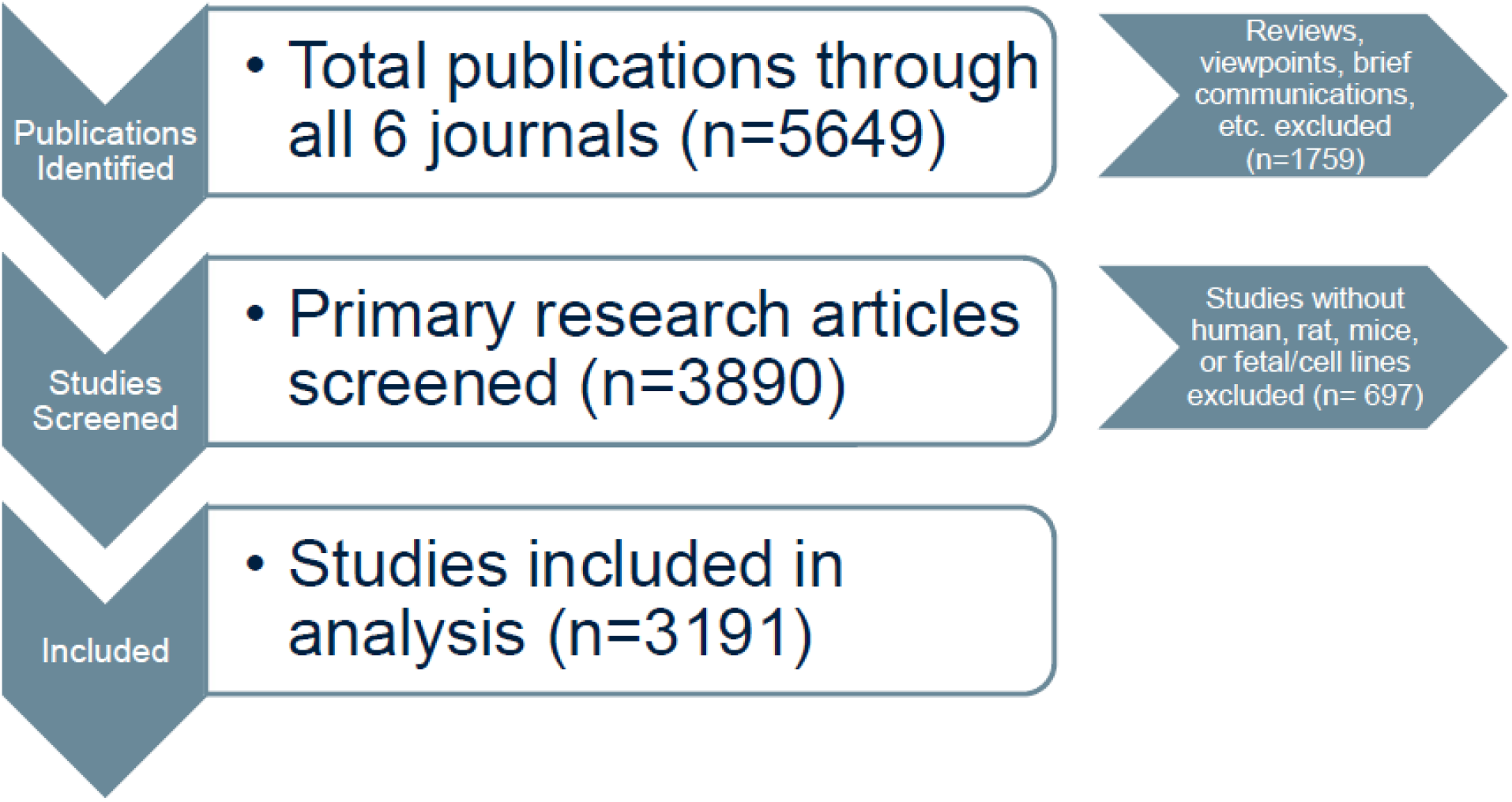
Inclusion of studies from all 6 journals. Reviews, viewpoints, brief communications and any other non- primary research articles were excluded. A total of 2456 studies did not match the inclusion criteria and were excluded. Only primary research articles containing human, rat, mice or fetal/cell lines were analyzed further (n=3191).

### Categorization of Inclusion of males and females and Sex-Based Analyses

Studies that matched the inclusion criteria were first examined to determine whether they included males and females, males only, females only, did not report sex, or were inconsistent throughout (i.e. used males in one experiment, and both sexes in another). If the study looked at both sexes, we determined whether there was balanced design (an equal ratio of male to female subjects). An unbalanced design was defined as one sex accounting for more than 60% of the total sample size.

Studies that included both sexes were then examined to determine whether they included any form of analysis using sex as a factor. Studies that did any type of sex analysis were then broken down into six categories: main effect of sex only, complete analysis by sex, sex as a covariate, analyzed sexes separately, statistics not given, and “mixed analysis”. Studies that only tested for a main effect of sex (examining differences between males and females on the dependent variable of interest) without regard to whether there were any interactions with other independent variables or any other further analyses were classified as “main effect”. An interaction effect examines the effect of sex along with other independent variables (e.g. treatment, genotype, disease). A significant interaction will indicate that the effect on the dependent variable (e.g. neurogenesis) varied across two independent variables, such as neurogenesis levels would differ by drug treatment based on the sex of the subject. Studies which analyzed the main effects and interaction effects of sex were classified as “complete analysis by sex”. Studies that used sex a covariate effectively removes the linear association of the variable sex from the dependent variables of interest. A covariate is a way of eliminating the variability due to sex, not analyzing for sex, and in doing so covariates are often referred to as ‘nuisance’ or ‘confound’ variables. Some papers stated that there was or was not an effect of sex but provided no statistical evidence to back up the statement and these papers were classified as “statistics not given”. A “mixed analysis” category was also included which consisted of studies which were inconsistent in their analyses throughout the study (i.e. analyzed sex in one experiment but did not analyze by sex in subsequent experiments). Any studies that used both sexes but did not mention any effects or analyzes by sex and therefore did not fit into any of these “analyzed” categories were classified as “not analyzed”. When sex information and analyses were only reported in the supplementary section of the studies, these studies were put into a “supplementary only” category. When a study analyzed by using sex as a discovery variable this meant that sex was used as a predictor/between-subject variable in the analysis and analysed for main and interaction effects, we refer to this as an optimal analysis for possible discovery of sex differences.

When a study employed a relatively equal sample size of both males and females and used them consistently throughout the study we refer to this as an optimal design for discovery of sex differences. Our reasoning behind this is that because unequal sample sizes affect power (the chance that the study will detect a sex difference if a sex difference exists or rejecting the null hypothesis when it is false) and if unequal sample sizes are paired with heterogeneity of variance this will affect the robustness of parametric tests^27^. This underscores that relatively equal sample sizes are necessary for an optimal design for discovery of possible sex differences. Modelling of sample sizes needed for discovery of sex differences suggest that when an interaction is present (interaction is when factor A has a different effect dependent on sex), high power can exist depending on the effect size of the interaction. For example, using a factorial ANOVA, high power (i.e β> 0.8) is obtained with relatively small sample sizes (n=5 per group) when the interaction shows either a reverse effect between sexes or no effect in one sex versus the other^21,28^. Larger samples sizes are needed when an interaction exists due to half of the effect in one sex over the other sex (β> 0.8, n=25 per group)^21^. Indeed the use of sex as a discovery variable can lead to increased statistical power, particularly when there are interaction effects indicating the sexes show opposing effects of a treatment/intervention on the variable of interest^21,28^. Thus, it is important that researchers not just consider that sex differences will result in overall (main) effects but that they may result in interaction effects (when a treatment has different effects in one sex versus another).

We refer to these designs and analyses as optimal for discovery of possible sex differences, as it would be impossible to detect any sex differences if the data were not analyzed by sex and if the sexes were not used consistently or the sample size employed was not advantageous to the discovery of possible sex differences. By using the word optimal we do not mean to imply that the studies were not optimal in the design for the particular experiment but that the design or analyses were not optimal for the discovery of any possible sex differences.

The country/region of origin of each paper was also examined. We included six categories for the region: USA, Canada, Europe (EU), the United Kingdom (UK), Asia (all countries in the continent of Asia), and combination/other. Combination/other refers to studies done by researchers from multiple countries, or from a country/region other than those previously mentioned.

We also examined the sex/gender of the first and last author of each paper. We determined author sex/gender by searching for the author online and looking for descriptions of them. When this was not possible we used the website genderize.io, a database which determines the sex/gender of a first name and provides a certainty factor associated with the name.

### Statistical Analyses

As the number of papers published differed by journal and year (Table 1) from a low of 55 (2019, *Molecular Psychiatry*) to a high of 1067 (2009, *Journal of Neuroscience*), we used proportional variables within each analysis. Data were reported and analyzed as percentages of total papers per journal per year. We used proportional data to run general linear analysis of variance (ANOVA) across year (2009, 2019) and discipline (Neuroscience, Psychiatry) with our dependent variables of interest. We also used method of analyses (complete analysis by sex, covariate, main effect, statistics not given, analyzed separately, mixed), single sex studies (male, female) and country of origin (USA, Canada, UK, EU, Asia, Combo) as within-subjects factors. Post-hoc comparisons used Newman-Keuls comparisons. Significance was set at α=0.05 and effect sizes are provided. Effect sizes using n_p_^2^ or Cohen’s d are provided. All analyses were tested for assumptions of ANOVA using Bartlett’s test of homogeneity of variance and Kolmogorov-Smirnov test for normality. None of the variables violated assumptions except for male only papers and these data were transformed prior to analysis.

**Table 1.** The number of papers examined that were published in 2009 or 2019 in the six journals investigated.

| Journal | Number of Papers |
| --- | --- |
| Neuron 2009 | 159 |
| Neuron 2019 | 207 |
| Nature Neuroscience 2009 | 118 |
| Nature Neuroscience 2019 | 143 |
| Journal of Neuroscience 2009 | 1067 |
| Journal of Neuroscience 2019 | 588 |
| Molecular Psychiatry 2009 | 70 |
| Molecular Psychiatry 2019 | 55 |
| Biological Psychiatry 2009 | 245 |
| Biological Psychiatry 2019 | 136 |
| Neuropsychopharmacology 2009 | 209 |
| Neuropsychopharmacology 2019 | 196 |
| **Total** | **3193** |

### Data Availability

Source data analyzed during the current study are available in the Dataverse repository, https://doi.org/10.5683/SP3/VDH895.

## RESULTS

### Most Neuroscience papers used rodents, whereas most Psychiatry papers used human subjects

We categorized the papers reviewed by subject species or tissue (Figure 2). Although the majority of studies in Psychiatry journals used human subjects this was closely followed by rodent studies whereas the majority of studies in Neuroscience journals used rodents which was three times higher than studies using human subjects. Neuroscience published three times more studies using cell lines than Psychiatry.

**Figure 2.**
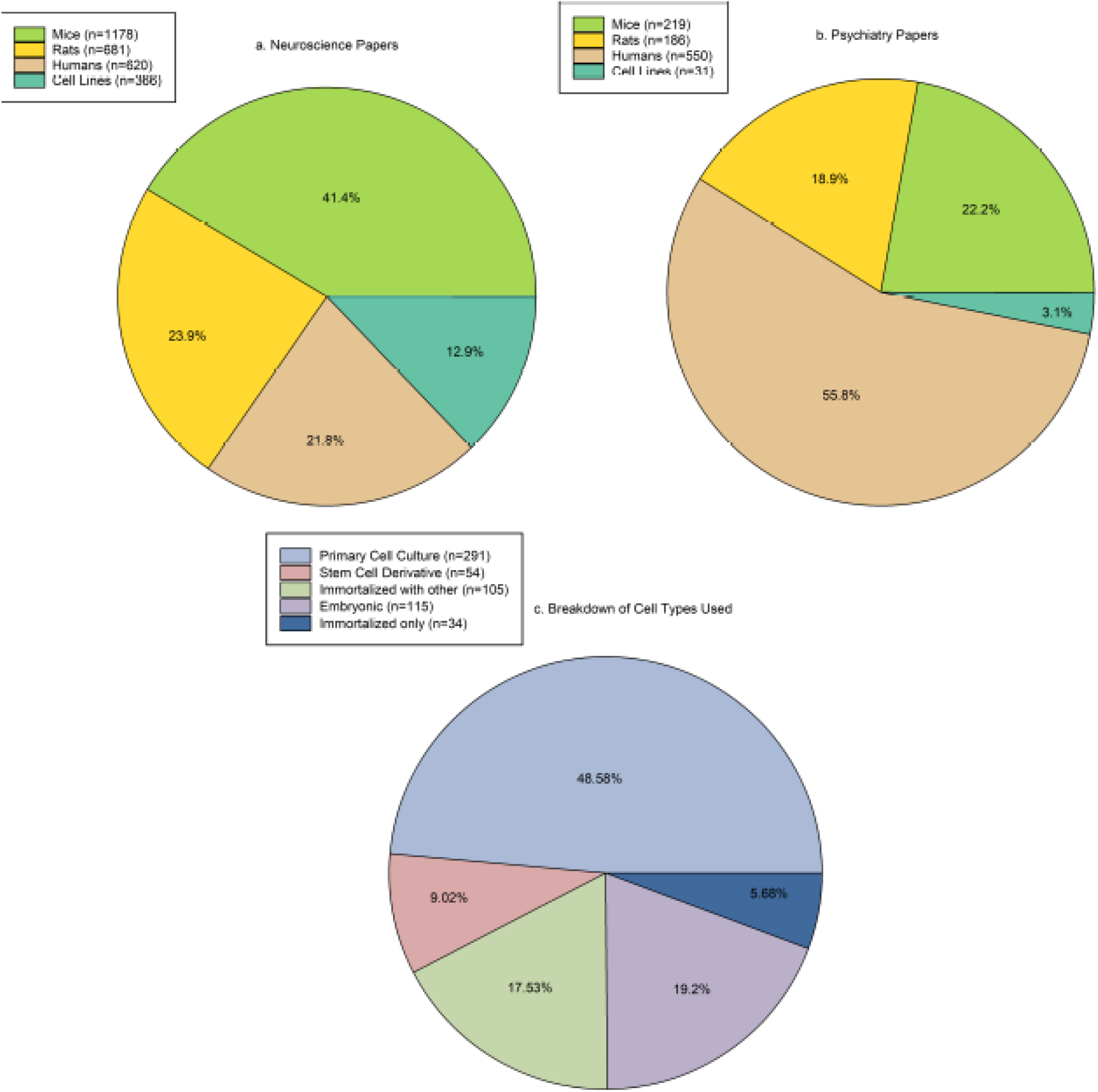
Reported species model used from each study. (a) Rodents (mice (n=1178) and rats (n=681)) were the most common species by studies in the Neuroscience discipline. (b) Human subjects (n=550) were the most common species used in Psychiatry studies. n=sample size Sample sizes are the number of papers that used the model systems and will total to greater than 3191 as some studies used two or more model systems. Of the studies that used cell lines, the majority used primary cell culture (n=291). The other types of cell line used were stem cell derivatives (n=54), immortalized with other cell types (n=105), embryonic (n=115), and immortalized only (n=34). Sample sizes will add up to greater than 397 as some studies used two or more cell lines in their papers. (c) Breakdown of type of cell line used. The largest proportion of studies used primary cell lines. Sample sizes are the number of papers and will add up to greater than 397 as some studies used two or more cell lines in their papers. We relied on the paper to distinguish whether cell lines were conducted in males or females, regardless of the cell line used.

### Neuroscience papers including males and females doubled from 2009 to 2019

Each paper was examined to determine whether any part of the paper mentioned the use of both sexes in the study, even if the data were not shown. Across all years and disciplines, the majority of all papers mentioned using both sexes (52.93%), which increased by 30% over the ten years (to 68.01% in 2019). Overall, just less than half (45.28%, n=962) of all Neuroscience publications mentioned using both sexes, while 60.58% (n=377) of all Psychiatry publications mentioned using both sexes. Neuroscience publications using both sexes significantly increased over the ten years by 50% to an astonishing 70.39% in 2019 (*p*=0.003; Cohen’s d=9.154). Psychiatry publications increased by just 10% to 65% in 2019 (*p*=0.32; interaction effect of year by discipline: F(1,8)=8.844, *p*=0.017, n_p_^2^= 0.525; Figure 3a). There were also significant main effects of year (F(1,8)=20.02, *p*=0.002, n_p_^2^= 0.714) and discipline (F(1,8)=5.14, *p*=0.05, n_p_^2^= 0.39).

**Figure 3.**
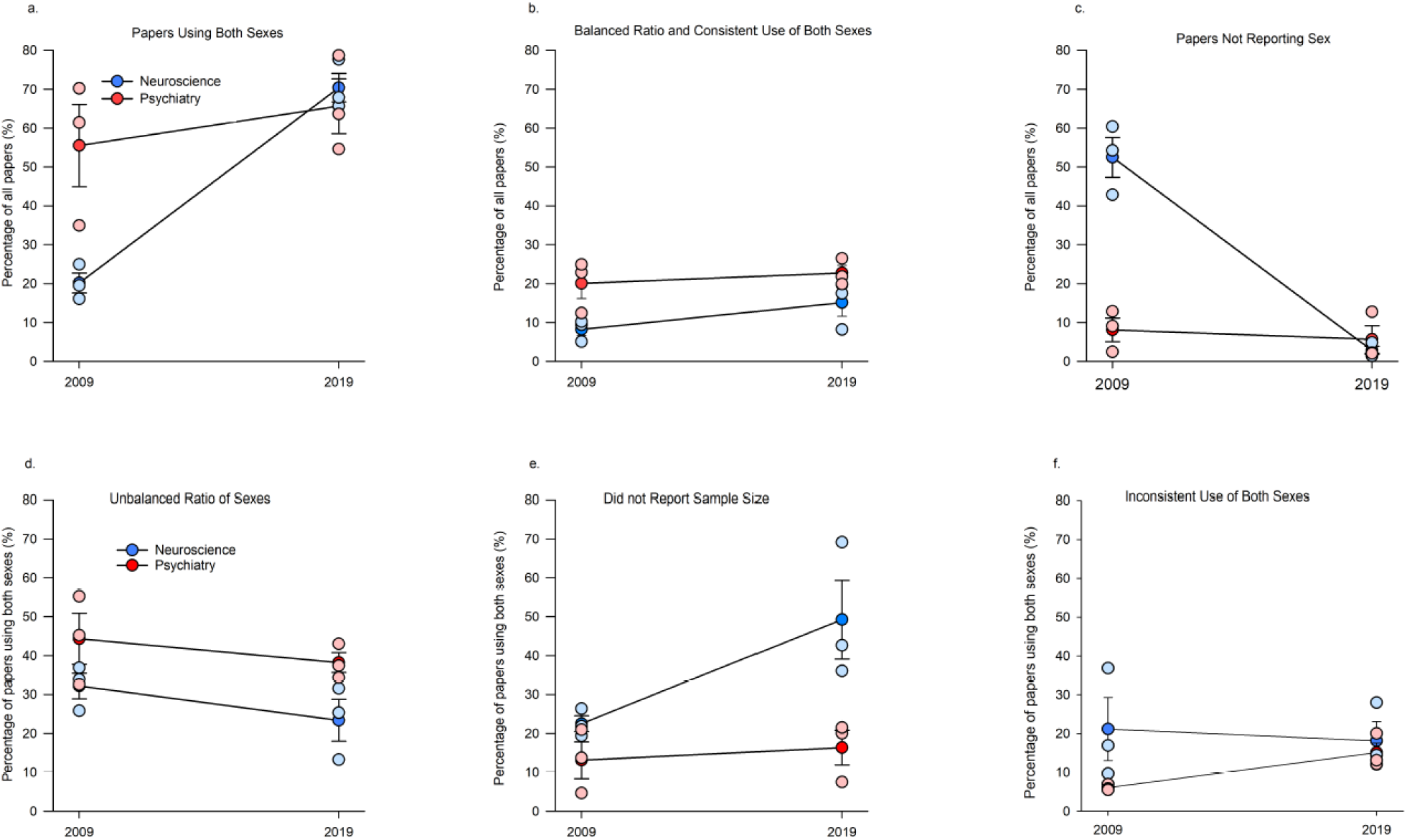
The percentage of papers including both sexes (a-c) and the breakdown of how studies that reported using both sexes are using them (d-f). (a) Percentage of papers using both sexes in any aspect of the study, regardless of consistency or balanced ratios. The percentage of papers including males and females increased significantly for Neuroscience p=0.003 but not Psychiatry papers (p=0.32). Number of papers: Neuroscience 2009 n=316, 2019 n=646; Psychiatry 2009 n=288, 2019 n=249. (b) Percentage of papers using both sexes consistently throughout the study with balanced ratios of the sexes. Number of papers: Neuroscience 2009 n=130, 2019 n=158; Psychiatry 2009 n=103, 2019 n=87. (c) Percentage of papers not reporting sex (sex omission) was decreased in the Neuroscience discipline p<0.001). Number of papers: Neuroscience 2009 n=617, 2019 n=25; Psychiatry 2009 n=34, 2019 n=14.(d) Unbalanced design (i.e. more than 60% of the subjects were one sex) was 34.52% of all papers including both sexes (number of papers: Neuroscience 2009 n=105, 2019 n=154; Psychiatry 2009 n=142, 2019 n=98) (e) Papers using both sexes but not disclosing sample sizes, are increasing in Neuroscience papers but not Psychiatry papers. (number of papers: Neuroscience 2009 n=69, 2019 n=304; Psychiatry 2009 n=27, 2019 n=38). (f) Inconsistent use of sex (i.e. using a balanced ratio in one aspect of the design, and an unbalanced ratio or one sex only in another aspect) accounted for 15.11% of studies that used males and females (number of papers: Neuroscience 2009 n=55, 2019 n=102; Psychiatry 2009 n=17, 2019 n=34). Means ∓ standard error of the mean.

However, the papers that included males and females included studies that mentioned the inclusion of both sexes but did not show these data. We then calculated a more rigorous count of the inclusion of sexes by including only studies that examined sexes in a balanced design and consistently used males and females throughout all the experiments in the paper. This more stringent criteria of inclusion of both sexes, resulted in a drop to below 20% of studies that used sex as an optimal design for discovery of possible sex differences (16.54% overall, 14.15% in 2009 to 18.93% in 2019). Psychiatry publications were twice more likely to use both sexes compared to Neuroscience publications (main effect of discipline, (F(1,8)=11.19, *p*=0.01, n_p_^2^= 0.583). There was no main effect of year (F(1,8)=2.715, *p*=0.137, n_p_^2^= 0.253) or interaction (F(1,8)=0.532, *p*=0.48, n_p_^2^= 0.062; Figure 3b).

The percentage of papers failing to disclose sex fell dramatically over the years, with the greatest change seen in Neuroscience as only 3% of papers omitted sex in 2019 (p<0.0001, Cohen’s d=7.73) as there was no significant change in Psychiatry papers across the years (p=0.63, Cohen’s d=0.43); discipline by year interaction (F(1,8)=45.21, *p*<0.001, n_p_^2^= 0.849, Figure 3c). There were also main effects (discipline: (F(1,8)=34.97, *p*<0.001, n_p_^2^= 0.813; year: (F(1,8)=55.2, *p*<0.001, n_p_^2^= 0.873).

### Most studies did not use an optimal design to discover sex differences

Although the percentage of studies using both sexes has increased, there are changes in the way that sex is being reported or used. What is driving the large discrepancy between the majority of all studies using both sexes but less than 20% of studies using sex optimally for discovery of possible sex differences? There were several scenarios we encountered in studies that used males and females which included 1) sample sizes were not given (25%), 2) the proportions of the sexes were dramatically different (34%), or 3) the use of sex was not used consistently throughout the studies (15%, Figure 3d-f).

Of the papers that used both sexes, just over a third of studies did not use a balanced design, with more Psychiatry papers employing this practice (main effect of discipline: F(1,8)=8.189, *p*=0.021, n_p_^2^= 0.505, Figure 3d). There were no other effects (all p’s > 0.153).

Just over a quarter of the papers that used both sexes did not identify sample sizes, which has effectively doubled across the years to almost a third of all studies that used both sexes in 2019 (32.79%) and this practice is twice as high in Neuroscience (35.87%) compared to Psychiatry (14.71%; Figure 3e; main effects: year (F(1,8)=6.06, *p*=0.039, n_p_^2^= 0.431) discipline: (F(1,8)=12.08, *p*=0.008, n_p_^2^= 0.602). Inspection of the graph indicates the increase across years is driven by Neuroscience as the percentage more than doubled in 2019 (49.25% from 22.49%; *a priori p*=0.014, Cohen’s d=2.16), whereas the percentage did not significantly change across the ten years in Psychiatry (from 13.09% to 16.32%, *p*=0.72; interaction (F(1,8)=3.73, *p*=0.089, n_p_^2^= 0.318).

The percentage of inconsistent use of sexes across the studies within a paper was 15.11% of all those that indicated they used both sexes. This percentage did not significantly change by year or by discipline (Figure 3f; *p*’s >0.10).

Few (4%) papers referred to the sex effects in the supplemental section and there were no significant differences across year or discipline (*p*’s >0.28; Table 2).

**Table 2.** The proportional percent of times that male and female data was found in the supplemental section. There were no significant differences by year or discipline. Overall four percent of papers referred to data on males and females in the supplemental section, not in the main body of the paper. n=Number of papers.

| Discipline | Mean ± SEM 2009 | Mean ± SEM 2019 |
| --- | --- | --- |
| Neuroscience | 3.9 ± 2.0 (n=3) | 4.2 ± 4.2 (n=14) |
| Psychiatry | 1.2 ± 1.2 (n=6) | 6.9 ± 2.1 (n=16) |

### Male-only papers disproportionally outnumbered female-only papers

Male-only papers were 9 times more common than female-only papers, regardless of year (main effect of sex: F(1,8)=324.39, *p*<0.0001, n_p_^2^= 0.976; Figure 4a). The percentage of studies that only included one sex remained constant across years (27% in males, 3% in females; *p*=0.36, Cohen’s d=0.0359) and did not differ across disciplines (*p*=0.34, Cohen’s d=0.932).

**Figure 4.**
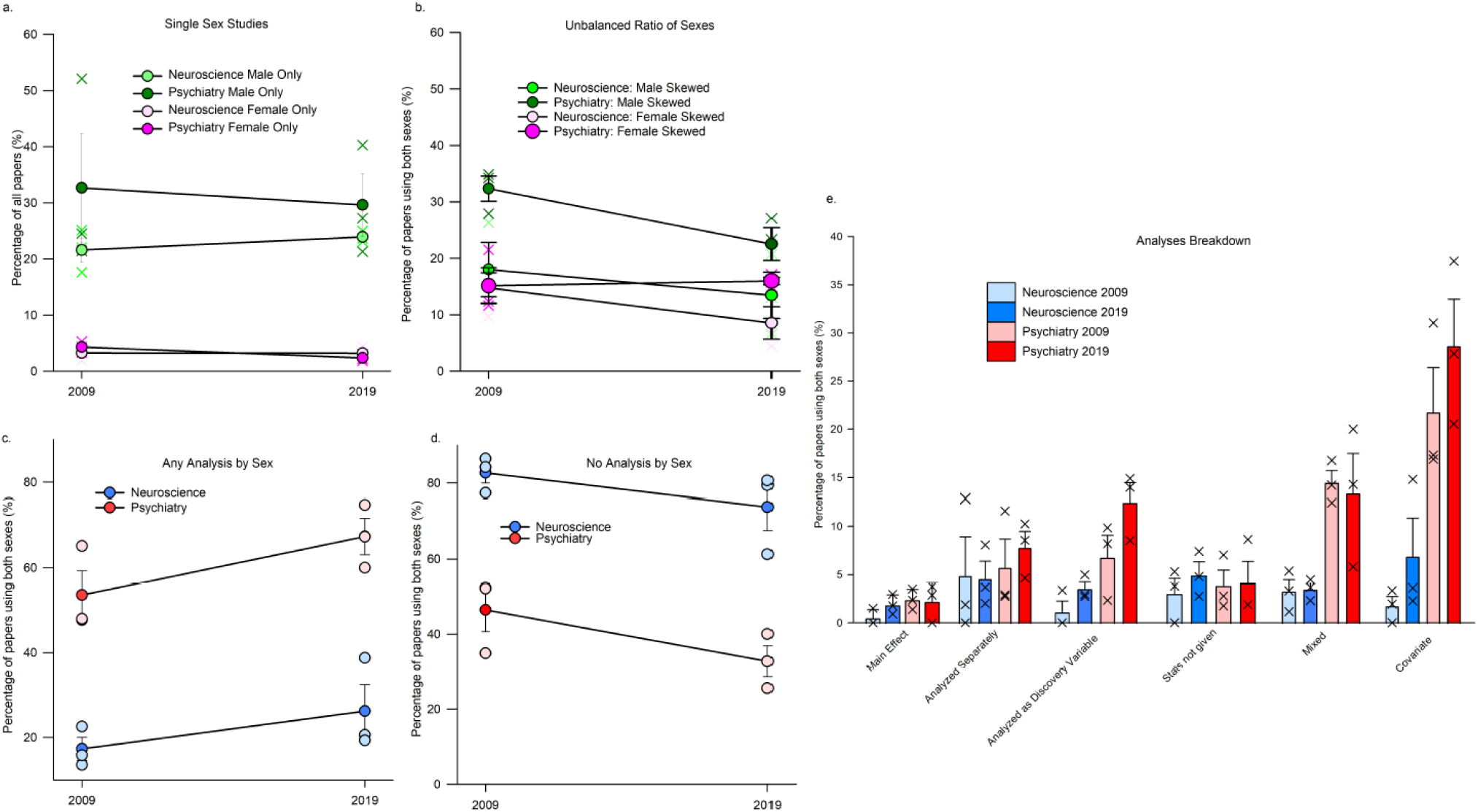
(a) Percentage of single sex studies across years and disciplines. Male only studies (26.96%) were 8.4 times higher than female only (3.29%) studies (number of papers: Male only: Neuroscience 2009 n=322, 2019 n=229; Psychiatry 2009 n=184, 2019 n=123; Female only Neuroscience 2009 n=55, 2019 n=35; Psychiatry 2009 n=23, 2019 n=10). (b) Of the studies using an unbalanced ratio of sex, there were more studies with greater proportion of males compared to females (number of papers: Male skew: Neuroscience 2009 n=56, 2019 n=84; Psychiatry 2009 n=97, 2019 n=60; Female skew Neuroscience 2009 n=56, 2019 n=70; Psychiatry 2009 n=51, 2019 n=39). As the percentage is proportionally based on the number of publications that year per journal the number of papers will vary differently that the proportional representation. (c) Breakdown of the type of analyses used by papers that used both sexes. Categories of sex analysis include: main effect of sex, sexes analyzed separately, sex analyzed as a discovery variable, stats not given (i.e. state some analysis was done but did not provide any statistics) mixed (i.e. any combination of analyses which may or may not be consistent throughout the study), and sex as a covariate. Number of papers: main effect: Neuroscience 2009 n=4, 2019 n=12; Psychiatry 2009 n=8, 2019 n=5; sex analyzed separately: Neuroscience 2009 n=9, 2019 n=22; Psychiatry 2009 n=12, 2019 n=19; analyzed as discovery: Neuroscience 2009 n=9, 2019 n=27; Psychiatry 2009 n=24, 2019 n=34; stats not given: Neuroscience 2009 n=11, 2019 n=32; Psychiatry 2009 n=8, 2019 n=7; mixed: Neuroscience 2009 n=5, 2019 n=19; Psychiatry 2009 n=39, 2019 n=38; covariate: Neuroscience 2009 n=6, 2019 n=33; Psychiatry 2009 n=54, 2019 n=75. (d) Majority of papers using both sexes did not analyze by sex, but this decreased slightly over 10 years. Number of papers: Neuroscience 2009 n=270, 2019 n=498; Psychiatry 2009 n=143, 2019 n=76. (e) Any analysis of sex in studies using both sexes. Psychiatry papers were more likely to perform any type of sex analysis than neuroscience papers. Neuroscience 2009 n=46, 2019 n=148; Psychiatry 2009 n=145, 2019 n=173. Means ∓ standard error of the mean.

Of the papers that used males and females in an unbalanced design, almost twice more were skewed towards males (main effect of sex skew: F(1,8)=20.23, *p*=0.002, n_p_^2^= 0.717) and there were almost double the percentage of sex-skewed papers in Psychiatry journals compared to Neuroscience (main effect of discipline F(1,8)=9.017, *p*=0.017, n_p_^2^= 0.531). There were no other effects (*p*’s >0.121); Figure 4b).

A thematic analysis on the responses that were given as to why single sex studies were used revealed of 51 documented responses, most referenced the need to reduce variability or confounds (50.98%, Table 3).

**Table 3.**
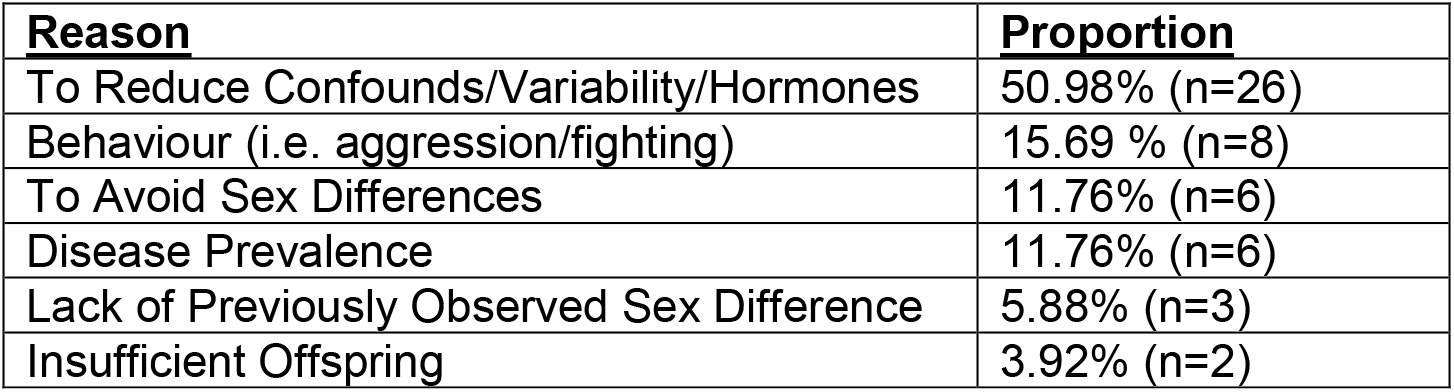
Thematic themes given for only including one sex in study design. 50.98% of single sex studies that gave a reason for the use of one sex referred to the reason to reduce confounds or variability mainly due to fluctuating hormones. N=number of papers

| Reason | Proportion |
| --- | --- |
| To Reduce Confounds/Variability/Hormones | 50.98% (n=26) |
| Behaviour (i.e. aggression/fighting) | 15.69 % (n=8) |
| To Avoid Sex Differences | 11.76% (n=6) |
| Disease Prevalence | 11.76% (n=6) |
| Lack of Previously Observed Sex Difference | 5.88% (n=3) |
| Insufficient Offspring | 3.92% (n=2) |

### The majority of papers did not analyze by sex

Of the studies that indicated they used both sexes, 40.34% said they analyzed their data by sex. The percentage of papers that indicated they did an analysis by sex increased from by 10% to 46.36% in 2019 irrespective of discipline (main effect of year: F(1,8)=5.17, *p*=0.05, n_p_^2^= 0.39). However, Psychiatry papers were three times more likely to have analyzed by sex compared to Neuroscience papers (main effect of discipline: F(1,8)=60.27, *p*<0.0001, n_p_^2^= 0.88). There was no significant interaction (*p*=0.79; Figure 4c).

Overall, whereas the majority of papers indicated they used both sexes, the majority of these studies did not analyze by sex (58.89%). Neuroscience papers using both sexes were almost twice more likely to not analyze by sex (78.24%) compared to Psychiatry papers using both sexes (39.53%; F(1,8)= 61.01, *p*<0.001, n_p_^2^= 0.884). Over the years the percentage of papers not analyzing by sex has significantly decreased by just over 10%, but unfortunately still remains above 50% (F(1,8)=5.24, *p*=0.05, n_p_^2^= 0.3955; Figure 4d).

### Only 6% of papers that used both sexes analyzed by sex

We further broke down how the papers analyzed by sex into 6 categories: complete analysis by sex (analyzed as a discovery variable), stats not given, covariate, main effect, analyzed separately, and mixed. Of the papers that used both sexes, 6.00% used sex as a discovery variable. Of the studies that used both sexes, the largest percentage of studies used sex as a covariate (14.36%) (Figure 4e).

Psychiatry papers were 5 times more likely to analyze using sex as a covariate (*p*=0.0001, Cohen’s d=2.998) or a mixed analyses (*p*=0.003, Cohen’s d=2.989) compared to Neuroscience papers, regardless of the year (Analyses Type by Discipline: F(5,40)=10.23, *p*<0.001 or *p*<0.0001, n_p_^2^= 0.56)). Covariate analyses were more often used than any other analysis (*p*’s< 0.001; main effect of Analysis Type: F(5,40)=13.14, *p*<0.0001, n_p_^2^= 0.62). There was also a main effect of Discipline (F(1,8)=60.27, *p*<0.0001, n_p_^2^= 0.88) and a main effect of Year with 2019 being higher than 2009 (F(1,8)=5.17, *p*=0.05, n_p_^2^= 0.39), but no other effects (*p*’s> 0.43,n_p_^2^ <0.11).

### North American papers increase use of sex in analyses

We next examined where papers originated. If we compared the total percentages of country origin to that percentage that used an optimal analysis for discovery of possible sex differences (Figure 5) one can see this increased for papers originating in the USA, Canada and a combination of countries, but fell for papers originating in the EU, Asia, and the UK. We did an analysis across years using the proportional data based on the number of publications that used both sexes by country (using each country as its own baseline). There were very low percentages across all countries with no significant difference across countries (p =0.39, n_p_^2^= 0.118) by year or discipline (p’s> 0.51, n_p_^2^=<0.10; Figure 5g-j). There were no other significant effects (p’s>0.05; number of papers Supplement Table 1).

**Figure 5.**
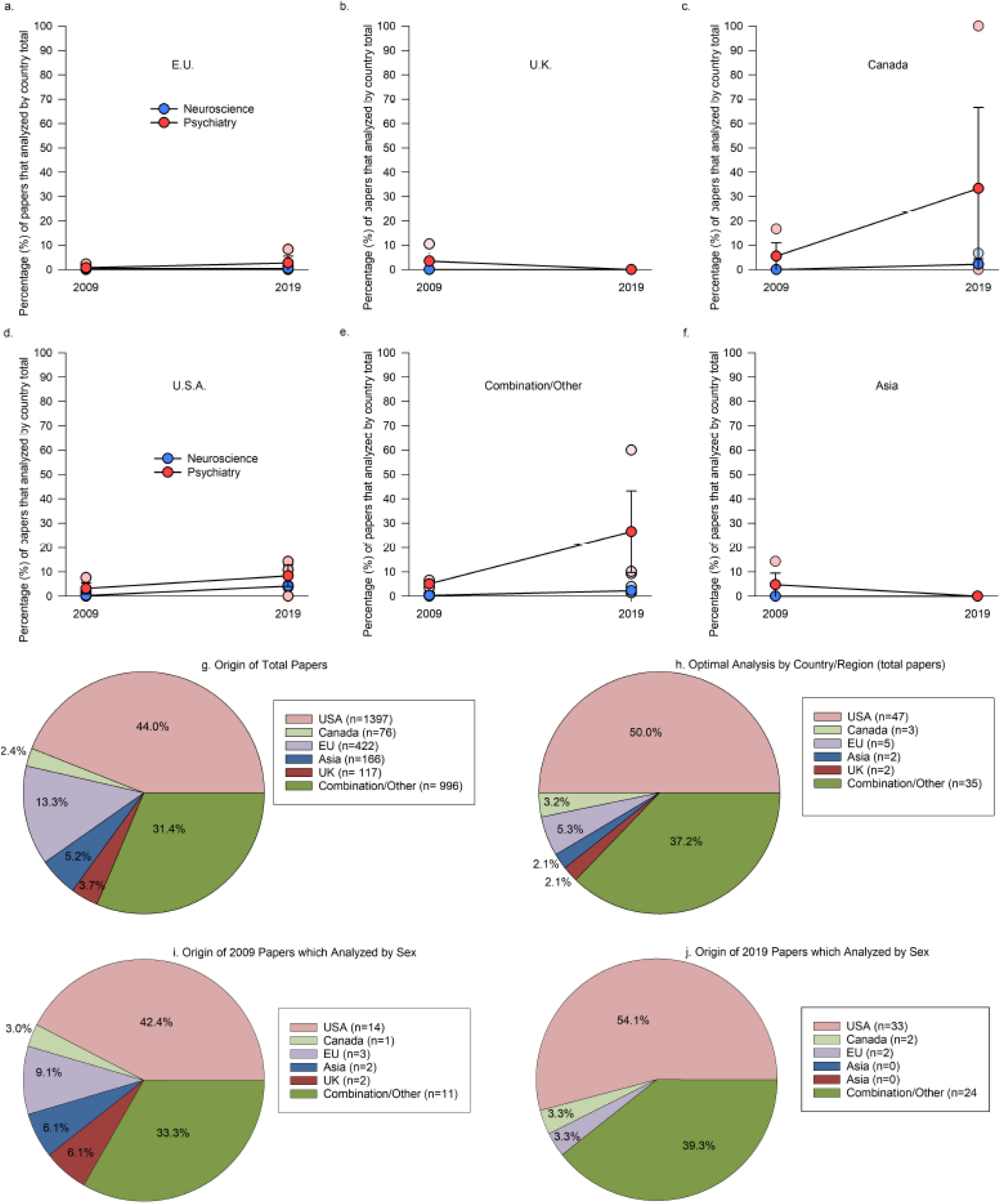
(a-f) Each country or combination of countries and their percentage of papers that analyzed using sex as a discovery variable across years compared to the country total. Papers originating from the USA, Canada, EU and a combination of countries had an increased percentage but none of these were significant. (a) E.U. is the European Union, (b) U.K. is the United Kingdom, (d) U.S.A. is the United States of America. (g) Country/region of origin of the paper and (h) breakdown of papers using optimal analysis for discovery of sex differences by region. (i-j) Origin of countries which analyzed using sex as a discovery variable compared to the total number of papers which analyzed by sex in 2009 (i) and 2019 (j)

### Females as first or last author increase analysis by sex

We examined whether sex/gender of the first or last author influenced the percentage of studies that used sex as a discovery variable, an optimal design for discovery of possible sex differences, or single sex papers. As these estimates are based on names we take a qualitative approach. Studies that listed males as first and last authors had a reduced (by 14%) proportion of papers that considered sex as a discovery variable compared to those that used an optimal design for the discovery of possible sex differences (Supplement Figure 2a-d). However, having a female as first author increased the percentage (market share) of papers that used sex as a discovery variable (increases of 4% with female/female and 10% for female/male). Comparing authorship sex/gender by male or female only papers (Supplement Figure 2c-d), shows a marginal increase for more female only papers when females are listed as the last author.

## Discussion

Our exhaustive survey of 3191 papers across six journals in Neuroscience and Psychiatry revealed some interesting insights into the inclusion, use, and analyses of both sexes in research over the ten-year period from 2009 to 2019 (Figure 6). Most studies used males and females in 2019, a 30% increase from 2009, irrespective of discipline. On the face of it, this is a dramatic positive benefit arising for greater knowledge and awareness on the importance of sex and gender as variables in research. However, the way researchers are reporting the use and analyses of males and females is not optimal for discovery of possible sex differences. This is troubling as collectively science will lose out on valuable information if researchers are neglecting to embrace the power of studying potential sex differences. When we determined the percentage of studies that used an optimal design for discovery of sex differences, the percentage of studies fell to 16.5%, a far cry from most papers that report the use of both males and females. Of the papers that reported using both sexes, three quarters of these papers either did not specify sample size, used unequal proportions of the sexes or used the sexes inconsistently within the paper. Perhaps even more concerning, most papers that describe using both sexes, did not analyze by sex (58%), and only 6% of studies used sex as a discovery variable across years and disciplines. Worse yet, the percentage of papers using optimal designs or analyses for discovery of sex differences has not meaningfully shifted in ten years across either discipline, despite the number of recent initiatives such as SABV, SGBA and SAGER. These findings should serve as a wake-up call to researchers, funders and journals, that if we are to harness the wealth of knowledge from studying both sexes, more needs to be done to improve the appropriate application of sex in reporting and analyses for discovery.

**Figure 6.**
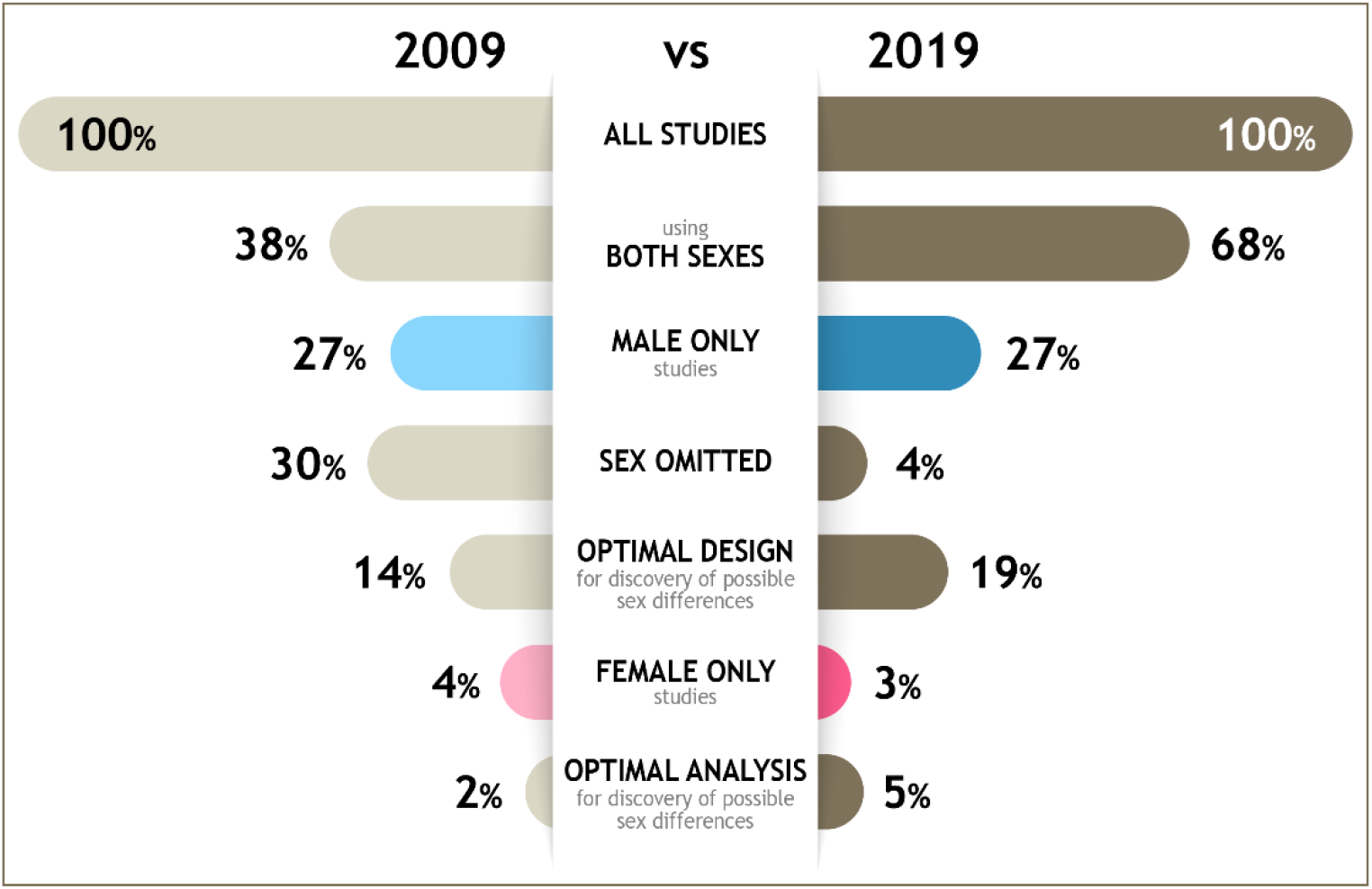
An infographic depicting the change in percentages of studies from 2009 and 2019 that used both sexes, single sex studies, studies that used an optimal design or analyses for the discovery of possible sex differences irrespective of discipline. Optimal design refers to relatively based sample size and use of males and females consistently across the experiments whereas optimal analyses refers to the use of sex as a discovery variable. Although the percentage of studies in Neuroscience and Psychiatry has increased dramatically the use of optimal design and analyses has not changed as dramatically and remain and low levels. There are nine times the percentage of male only compared to female only studies.

As noted, there has been a vast increase in the reporting of both sexes in both Psychiatry and Neuroscience papers from 2009 to 2019 to almost 70% in 2019. Neuroscience showed a 50% increase in reporting the use of both sexes over the years whereas the increase was 10% in Psychiatry over the same ten-year period. This difference is likely driven by the majority of papers using humans in Psychiatry journals, which may be a direct result of an earlier (2001) NIH mandate to include males and females in clinical research. The great majority of Neuroscience and Psychiatry articles are using both sexes in 2019, which is encouraging. Our finding of a 50% increase across ten years is also higher than the almost 20% increase seen from 2010-1014 in Neuroscience^25^ and the 34% increase seen by Woitowich and colleagues^24^ across the same ten year period. In addition, the 68% of studies that included males and females in 2019 in our study is notably higher than the 52% of Neuroscience papers reporting the use of both sexes in 2017^29^, likely reflecting an upward trend across years. The large progress made in Neuroscience across the 10 years was also noted by Woitowich^24^ who, as noted, saw an increase to 63% in 2019 using a sampling of 20 articles from 4 journals, two of which overlapped with ours (Journal of Neuroscience and Nature Neuroscience). In the present paper we exhaustively sampled from 3 journals in Neuroscience, much like the work by Meitzen and colleagues^25,29^ who exhaustively searched for Neuroscience papers in 6 journals, 3 of which overlapped with the journals we chose (Nature Neuroscience, Neuron, Journal of Neuroscience). Thus, collectively, multiple studies, using different journals and methods of sampling, consistently indicate that there is an increasing trend in articles that include males and females in their work.

Although the use of both males and females in research has been steadily increasing to include a majority of studies, research highlighting or mentioning sex differences is scarce. Why might this be? We examined whether papers were using optimal designs for discovery of possible sex differences. When we accounted for studies that did not disclose sample size of the sexes, used unbalanced design or only used both sexes in a portion of the study, we found that only 16% of studies used a design that was optimal for discovery of sex differences. Some researchers will argue that investigating both males and females is only important in the first step and thus the use of both sexes in further experiments, beyond the initial study is not required. However, there are numerous examples where a trait may not have sex differences but the mechanisms underlying that trait do show significant differences between males and females^30–33^. Thus, using males and females in one experiment does not preclude the fact that they may show differences in further experiments. Unfortunately, the use of the most advantageous design for discovery of sex differences was only employed in just under 20% of studies in 2019. Thus, although it appears on the face of it that most studies are using males and females, the majority of these studies do not incorporate sex in their design that is optimal for discovery of possible sex differences.

Our findings also demonstrated that 25% of studies using both males and females do not report sample size, consistent with the findings from Woitowich and colleagues^24^. Perhaps more concerning is that particularly in Neuroscience, this trend is increasing over the ten years with almost 50% of studies not reporting the sample size of males and females used. This trend is troubling as the reader is unable to judge how effectively males and females were used in the study.

As many other researchers have reported, most publications do not analyze by sex. Perhaps more concerning is that only 6% of studies that used males and females used sex as a discovery variable, which has not increased effectively over the years. This translates into only 4% of all publications examined that used sex as a discovery variable. Fourteen percent of papers that used males and females used sex as a covariate, with this statistical approach used more often in Psychiatry. A covariate removes the linear association of the factor of sex against the dependent variable, removing any linear variation due to sex. In our minds this is in opposition to the intention of SABV or SGBA. The point is not to remove the variation due to sex but to determine whether or not sex is a variable that could be causing differences in outcomes. Others have shown that the use of sex as a covariate can result in the reduction of power and the loss of important information when a sex difference is present^34^. Mersha and colleagues^34^ show that 26 more single nucleotide polymorphisms (SNPs) were identified in a sex stratified analysis compared to when sex was used as a covariate. Put another way, when sex was used as a discovery variable 47 SNPs were identified that were associated with asthma but if sex was used as a covariate only 21 SNPs were identified^34^. They also found that effect sizes were larger when a sex-stratified analysis was used, contrary to popular opinion that power would be negatively affected with the addition of sex as a discovery variable. Some argue that design and sample sizes are not powered to consider sex-stratified analyses, but if the sex effects are large, or in opposing directions, the resulting power with the inclusion of sex, will improve as others have demonstrated^21,28,34–36^. Taken together, our survey of the literature suggests that researchers are underestimating the power of using sex as a discovery variable in their research.

Similar to other reports in Neuroscience and other biological disciplines^23,25,28,29^, we found female only studies were a small percentage of studies. Our survey indicated that the percentage of female only studies is very low at 3%. Our findings are comparable with others showing that 5% of Neuroscience studies were female only in 2009^28^ and in 2017^29^. Although, the use of sex/gender in studies is important, single-sex studies are still needed. Given the dearth of information on women’s health and disparities in diagnosis^7^, and continued underrepresentation in clinical trials^13^, one could argue that we need female only studies even more so than male only studies - or that at least the single sex studies should be conducted and published in equivalent proportions. Indeed the impetus for SABV and SGBA was instigated in part because of the lack of knowledge of how females differed in their response to treatments and disease^37^. There are female-specific experiences that affect female health, such as menstruation, hormonal contraceptives, pregnancy and menopause that need to be studied^38–41^. Unfortunately, as highlighted by the current study, the percentage of studies that use only females is devastatingly low and has not improved over ten years. Funders and researchers should work to correct this imbalance.

The rationale for excluding females was often to “reduce variability”. To exclude females based on greater variation than males is not valid, as two studies have found that the variability between males and females is not different in rodents^42,43^. Although it is common to think that females will have more variability due to their hormones, males (rodents and primates) and females have diurnal fluctuations in cortisol/corticosterone^44,45^. Furthermore, human males have diurnal fluctuations in testosterone levels that vary significantly with age^44^. Researchers are encouraged to consider that many hormones that can vary with diet, age, housing conditions and experience across both sexes^46–48^. Thus, variability between males and females should not be a limiting factor in the use of males and females in research.

There have been calls in the literature to ensure that editors and reviewers of manuscripts ensure that published reports use both males and females and report on outcomes^49^. SAGER guidelines were developed by the European Association of Science Editors to improve sex and gender in research reporting in 2016^50^, and indeed, some journals have adopted SAGER guidelines including over 500 Elsevier journals^51^. Among the guidelines, it is recommended that authors include the sex in the title and abstract, background information on sex/gender effects on the variables of interest in the paper and in the results to disaggregate and analyze the data by sex/gender. However, the percentage of journals that have adopted SAGER are still low with one study finding under 10% of journals in Psychology had adopted the guidelines and in those journals the guidelines were only adopted for the title, abstract and methods but not on reporting of analyses or data by sex/gender^52^. However, as can be seen from the present data, the publishing of this information, particularly with respect to the analyses of sex as a discovery variable is limited, and a more concerted effort needs to be adopted.

We only examined three journals for each of the two disciplines, however we did an exhaustive search of eligible research papers within each journal, culminating in over 3000 articles reviewed. Contrast this to other papers that surveyed 841 articles across 2 years^24^ to over 6000 articles across 4 years^25^. We, as others^25^, selected journals based on ranking by ISI, with some overlap in journals chosen. However, our exhaustive search of these 6 journals gave values that were not appreciably different from those that used fewer papers within more journals, or exhaustive searching in a greater number of years, suggests either survey method yields similar results. Often the terms sex and gender were used incorrectly. Others have shown that in the fish literature, gender was used incorrectly 99% of the time^53^. Often gender is conflated with gender identity, and it is important to understand that gendered effects can be realised when considering a number of intersectional variables with sex/gender identity^54^. A final consideration is that for biomedical research at NIH, the SABV consideration was instituted in 2016 and this may not have given enough time to fully realise the potential in 2019 survey of the literature. However, the fact that in the Neuroscience journals there was a dramatic increase in the percentage of studies using both males and females to 70% suggests that there is some movement for inclusion, but this is unfortunately not transferring to analyses by sex.

### Call to Action: Fixing Implementation Issues with Carrots and Sticks

Given that there is excellent uptake in the use of both males and females in research, what is driving the lack of optimal design and analyses for discovery of sex differences? It seems possible that researchers themselves are not aware that they are not using best practices, perhaps due to the lack of consensus on how to use sex in analyses and the required sample size in the literature^37^. Three-quarters of researchers say they report the sex in their papers^55^, which matches our data. Of these researchers, 50% of them said they analysed findings by sex^55^ and our results show although that 40% of researchers analysed by sex in some fashion only 6% used sex as a discovery variable. Taken together, these data indicate that researchers may be considering analyses that are suboptimal or not reporting analyses even when they have done them. Another concerning factor, was that while researchers indicated they had a good knowledge of SABV they incorrectly used sex and gender in discussing their views, indicating a lack of knowledge. Thus, it is possible that researchers believe that the addition of both sexes without thorough analyses is enough to satisfy the initiatives. Another outcome of the qualitative analyses, that perhaps should not come as a surprise, is that some researchers do not appreciate mandates^56^.

One could argue that the mandates do not go far enough and are limited to a few agencies in the EU, Canada, and the US. There are also no repercussions when authors do not publish or analyse by sex. Indeed, NIH funding did not significantly affect the percentage of studies that analysed by sex (included covariate) with a net increase of just 3% (to 9%) overall^29^. Our data indicate that there is a non-significant increase in studies that used sex as a discovery variable in the US, Canada, and the EU pointing to an overall benefit of the current mandates that exist in those countries. However, it is important to underscore that these were still low percentages and that there are no reporting requirements from these funding agencies.

What can funders do to promote more work on sex differences? One solution is to have funding dedicated specifically for SABV and SGBA proposals. Evidence suggests that this approach has been successful in cardiovascular research. The American Heart Association (US) has dedicated funding for sex differences and as a result sex and gender based research and analyses in cardiovascular disease has flourished^57^. To undertake a seismic shift, funders would make these funds a significant portion of the budget to provide enough incentive to entice researchers to think deeply about incorporation of sex in research. Dedicated funding would not only generate proposals and knowledge dedicated to the analyses of sex differences, but they would also have the by-product of creating the next generation of researchers that integrate sex into their research. One can also look at how significant funding to Amyotrophic lateral sclerosis (ALS) and AIDS advanced research in these areas. In 2014, the ice bucket challenge raised greater than $115M in the US and this attention leveraged dedicated funding from other sources tripling ALS research budgets in 5 years^58^. This bolus of funding doubled the number of ALS publications, led to a 50% increase in investigators interested in ALS, and has dramatically accelerated the number of clinical trials in ALS^59^. Scientific evidence takes time to build, but fruits of discovery with the increased funding are paying off with promising new treatments^60^. It’s hard not to get excited about the possibilities if this type of funding is extended to fill the sex disparities in health research. AIDS research is another success story with dramatic advancements in AIDS research that came with dedicated funding. AIDS research funding increased dramatically over the years to >18B^61^. With these dedicated funds have come advancements in therapeutics such that individuals with HIV can live relatively full lives^62^. To make significant progress, funders need to have dedicated funding for SABV/SGBA which would have a cascading effect to get more researchers interested in SABV/SGBA, ensure consideration of sex/gender as a discovery variable, increase the number of discoveries and train the next generation of SABV/SGBA researchers.

What can publishers do to promote publications using sex-based analyses? When journals adopt SAGER guidelines, it is up to the authors, reviewers and editors to ensure the guidelines are met. In over a third of submissions to a neuroendocrinology journal, authors and reviewers failed to notice that sex/gender had not been disclosed^63^. This suggests, not surprisingly, that not every researcher is triggered to think about the consideration of sex in experimental design and analyses. Training modules will help, but working on a similar premise as above, enticing researchers to explore the influence of sex/gender in their data may be a more fruitful approach. If journals, especially those with higher visibility, adopt calls for papers using sex and gender-based analyses this will serve as a catalyst to ensure more researchers consider possible sex differences and further promote the notion that this research is important to publish.

Lastly, our data are suggestive that countries that have adopted mandates for inclusion of sexes in research have a small positive effect to increase analyses by sex as a discovery variable. Compared to the country norms, papers originating from the US, Canada or a combination of countries had greater representation in using sex as a discovery variable. In addition, female first or last authors increases the use and analyses of both sexes in research^64,65^ which is consistent with our own data. Recently, there has been concerted efforts to promote diversity in science^66^ and these findings suggest that increasing sex/gender diversity in science is another fruitful pathway to improve the percentage of studies exploring sex as a discovery factor in analyses.

We hope these data are a wakeup call to the research community to not only include males and females in their research but to ensure appropriate methods of integration and analyses. If researchers are merely sprinkling in a few animals of the opposite sex in one of many experiments this will not allow for discovery of the impact of sex as a biological variable. Nor will the non-robust adoption of sex in experiments harness the additional power that the analyses of sex can afford^34^. Research shows us that the use of sex as a discovery variable can lead to fruitful knowledge, and can conclude that the different mechanisms between males and females require distinct treatment^21^. Indeed, inclusion of sex in analyses and design will improve not only the health of females but of males^67^. We lose collectively, not just in knowledge gained, but also in our search of more effective treatments when sex is not considered in the design and analyses of our studies. We call on funders, reviewers and researchers to recognize that sex and gender matter across all disciplines. The community needs to be aware that there are many types of sex differences^19,68^ and that some sex differences are revealed due to perturbations in environment, genotype, or disease^19,69,70^ so it is important to continually examine and analyze both sexes throughout the studies. It is imperative that more attention is paid to the appropriate design and analyses of sex/gender in the literature. We need to study how mandates can improve adherence in both study design and dissemination. To ensure precision medicine, we need the community of funders, researchers and publishers to embrace the addition of SABV, SGBA and SAGER to improve the health of women, men and gender-diverse individuals.

## Acknowledgements

This research was funded by a Natural Sciences and Engineering Research Council of Canada grant to LAMG (2018-04301) and supported by the BC Women’s Foundation.

## Author Contributions

RKR and TFLS collated and tabulated the data with consultation with LAMG and AA. TEH, AA, and LAMG carried out the analysis. RKR, TFLS and LAMG wrote the manuscript. LAMG supervised the work. All authors declare no competing interests.

